# Genomic differentiation and intercontinental population structure of mosquito vectors *Culex pipiens pipiens* and *Culex pipiens molestus*

**DOI:** 10.1101/867887

**Authors:** Andrey A. Yurchenko, Reem A. Masri, Natalia V. Khrabrova, Anuarbek K. Sibataev, Megan L. Fritz, Maria V. Sharakhova

## Abstract

Understanding the population structure and mechanisms of taxa diversification is important for organisms responsible for the transmission of human diseases. Two vectors of West Nile virus, *Culex pipiens pipiens* and *Cx. p. molestus*, exhibit epidemiologically important behavioral and physiological differences, but the whole-genome divergence between them was unexplored. In this study, we re-sequenced and compared the whole genomes of 40 individual mosquitoes from four locations in Eurasia and North America: the Republic of Belarus, the Kyrgyz Republic, and the United States of America. Principal Component, ADMIXTURE, and neighbor joining analyses of the nuclear genomes identified two intercontinental, monophyletic clusters of *Cx. p. pipiens* and *Cx. p. molestus.* The third cluster, having a polyphyletic origin, was formed by *Cx. p. pipiens* and *Cx. quinquefasciatus* from the USA. The level of genomic differentiation between the subspecies was uniform along chromosomes. The ADMIXTURE analysis determined signatures of admixture in all *Cx. p. pipens* populations but not in *Cx. p. molestus* populations. Comparison of mitochondrial genomes among the specimens showed a paraphyletic origin of the major haplogroups between the subspecies but a monophyletic structure between the continents. Thus, our study identified that *Cx. p. molestus* and *Cx. p. pipiens* represent different evolutionary units with monophyletic origin that have undergone incipient ecological speciation.

## Introduction

The advent of genomics has provided new insights into population divergence between incipient taxa, changing our vision about the mechanisms of adaptation and speciation ^1^. Experimental data demonstrate highly heterogenous patterns of population differentiation in various groups of organisms ^2^. In addition to the classical model of allopatric speciation, when incipient taxa are isolated geographically ^3^, it becomes obvious that ecological speciation or the development of reproductive isolation between populations as a result of adaptation to different environments is feasible and common in nature ^4–6^. Understanding the population structure and the mechanisms of taxa diversification in the changing environment is extremely important if the studied organisms are related to the transmission of human diseases ^7^. Members of the *Culex pipiens* complex are globally distributed throughout Europe, Asia, America, Africa, and Australia and represent competent vectors of the lymphatic filariasis parasite and encephalitis viruses, including the widely spread West Nile virus ^8–11^. However, despite the fact that *Cx. pipiens* was the first mosquito species described by C. Linnaeus in his “Systema Naturae” ^12^, mosquitoes from the *Cx. pipiens* complex still represent “one of the major outstanding problems in mosquito taxonomy” because the members of the complex can mate and produce viable progeny in nature ^13,14^. Thus, the mechanisms of genetic differentiation in the members of this complex remain poorly understood.

The “Catalog of the Mosquitoes of the World” (Knight, 1978), maintained by the Walter Reed Biosystematics Unit at the Smithsonian Institution (http://www.wrbu.si.edu), recognizes the following species as members of the *Cx. pipiens* complex: *Cx. pipiens*, *Cx. quinquefasciatus*, *Cx. australicus*, and *Cx. globocoxitus*. Among these species, *Cx. pipiens* and *Cx. quinquefasciatus* spread globally in temperate and tropical/subtropical regions, respectively. Distribution of *Cx. australicus* and *Cx. globocoxitus* is restricted to Australia. In addition to these species, the *Cx. pipiens* complex includes a subspecies, *Cx. p. pallens*, found in Japan and the Far East of Eurasia, and two additional members, *Cx. p. pipiens* and *Cx. p. molestus*, that, according to this catalog, represent two physiological forms. Indeed, *Cx. p. pipiens* and *Cx. p. molestus* exhibit important physiological and ecological differences ^8,15^. *Cx. p. pipiens* mates in open spaces, feeds on birds, and requires a blood meal for oviposition. During the winter, females of *Cx. p. pipiens* undergo diapause. In contrast, *Cx. p. molestus* mates in confined spaces, feeds on mammals, can lay eggs without a blood meal, and females of this subspecies cannot enter diapause. *Cx. p. molestus* is also known as an underground mosquito because it invades basements, sewers, and underground railways. This fact explains why *Cx. p. molestus* can survive in a cold climate without the ability to enter diapause ^8,16^. Since *Cx. p. pipiens* has strong bias toward feeding on birds, while *Cx. p. molestus* readily bites humans and other mammals, hybrids between the two forms can act as a bridge vector between the bird reservoirs and susceptible mammalian hosts ^17–19^. Thus, understanding the abilities of *Cx. p. pipiens* and *Cx. p. molestus* to produce viable progeny in natural populations is of medical importance.

Historically, P. Forskal described *Cx. molestus* as a separate species from Egyptian specimens in 1775, but, later, this species was synonymized with *Cx. pipiens* ^13^. Following the concept of E. Mayr ^3^, P. Mattingly proposed considering the *Cx. pipiens* complex as a polytypic species; thus, the status of *Cx. molestus* was reduced to the subspecies *Cx. p. molestus* ^20,21^. Later, E. Vinogradova considered *Cx. p. pipiens* and *Cx. p. molestus* as ecophysiological forms because of the differences in their behavior ^16^. R. Harbach argued that *Cx. p. molestus* is a “phenotypic, ecological, and physiological variant” of *Cx. p. pipiens* and should not be considered a subspecies because of the absence of clear morphological differences and the presence of hybrids ^13^. However, a molecular analysis based on microsatellite markers determined that *Cx. p. pipiens* and *Cx. p. molestus* from different continents cluster together in a subspecies specific manner and more likely represent two separate taxonomic units or species with a unique evolutionary history ^18^. Similar results regarding the monophyletic origin of *Cx. p. pipiens* and *Cx. p. molestus* were obtained by the analysis of single nucleotide polymorphisms (SNPs) in the mitochondrial Cytochrome Oxidase I (COI) gene of European populations ^22^. These conclusions were strongly supported by another work using amplified fragment length polymorphism markers that identified discrete genomic differences between the subspecies and, thus, considered *Cx. p. pipiens* and *Cx. p. molestus* to be distinct evolutionary entities that are likely in the process of incipient speciation ^23^.

Sequencing the *Cx. quinquefasciatus* genome offered exciting opportunities for comparative genomic studies of the *Cx. pipiens* complex ^24^. Here, for the first time, we used whole-genome analysis of individual mosquitoes to investigate the level of genomic differentiation and population structures of *Cx. p. pipiens* and *Cx. p. molestus* from different continents. We compared sequences of 40 samples from field collected mosquitoes from Eurasia (Minsk, Republic of Belarus and Mailuu-suu, Kyrgyz Republic) and North America (Washington, D.C., USA) and two laboratory colonies of *Cx. p. pipiens* and *Cx. p. molestus* derived from Chicago, IL, USA. In our study, we follow the classical nomenclature that was proposed by Mattingly in 1951 and consider *Cx. p. pipiens* and *Cx. p. molestus* as subspecies ^20,21^.

## Results

### Mosquito collections

In this study, we resequenced whole genomes of 40 individual mosquitoes from 4 different locations—two from Eurasia and two from North America (Table 1, Fig. 1). All mosquitoes were collected from the urban environment. *Cx. p. molestus* mosquitoes were collected at the larval stage from an underground water source in two locations (the Kyrgyz Republic and the Republic of Belarus). *Cx. p. pipiens* were also collected at the larval stage in an open water reservoir in the Kyrgyz Republic, but at the adult stage in the basement of a multi-floor building in the Republic of Belarus. All field collected specimens from Eurasia were identified as *Cx. pipiens* by larval morphology and then tested to subspecies level using the COI assay, which is based on SNPs in a specific position ^25^. Mosquitoes from the Washington, D.C., USA samples were collected as egg rafts from two different urban environments—an underground parking lot and an open water reservoir. Both egg collections were used to establish mosquito colonies that were kept in the laboratory for several generations. At the larval stage, mosquitoes from both collections were identified morphologically as *Cx. pipiens.* However, molecular assays based on COI SNP ^25^ and polymorphism in the flanking region of a microsatellite locus ^26^ failed to identify them at the subspecies level. The latter method was considered unreliable for the diagnosis of *Cx. p. molestus* and *Cx. p. pipiens* in California ^27^. In addition, we performed a physiological assay and identified that mosquitoes from the parking lot colony were able to lay eggs without a blood meal. Mosquitoes from the open water collection needed a blood meal for egg development. Thus, we called our samples from Washington, D.C, USA *Cx. pipiens* autogenetic (WPA) and *Cx. pipiens* anautogenic (WPN). In addition, we used two colonies of *Cx. p. molestus* and *Cx. p. pipiens* that were derived from the Chicago area in the USA ^23^.

**Table 1.** Collection of the *Cx. pipiens* mosquitoes identified and used for sequencing

| Sample ID | Country | City, state | GPS coordinates | Habitat | Species/subspecies, stage, physiology | Identification method | Sample size |
| --- | --- | --- | --- | --- | --- | --- | --- |
| BP | Republic of Belarus | Minsk | 54.00, 27.00 | Basement, multi floor building | *Cx. p. pipiens*, adults | COI, <sup>25</sup> | 5 |
| BM | Republic of Belarus | Minsk | 54.00, 27.00 | Basement, multi floor building | *Cx. p. molestus*, larvae | COI, <sup>25</sup> | 5 |
| KP | Kyrgyz Republic | Mailuu-su II | 41.00, 72.00 | Aboveground open water pool | *Cx. p. pipiens*, larvae | COI, <sup>25</sup> | 5 |
| KM | Kyrgyz Republic | Mailuu-su | 41.00, 72.00 | Underground barrel | *Cx. p. molestus*, larvae | COI, <sup>25</sup> | 5 |
| CP | USA | Chicago, IL | N/A | colony | *Cx. p. pipiens*, larvae | <sup>23</sup> | 5 |
| CM | USA | Chicago, IL | N/A | colony | *Cx. p. molestus*, larvae | <sup>23</sup> | 5 |
| WPN | USA | Washington, DC | 38.25, -76.76 | Aboveground puddle | *Cx. pipiens*, larvae, anautogenic | morphology | 5 |
| WPA | USA | Washington, DC | 38.25, -76.76 | Underground parking lot | *Cx. pipiens*, larvae, autogenic | morphology | 5 |

**Fig. 1.**
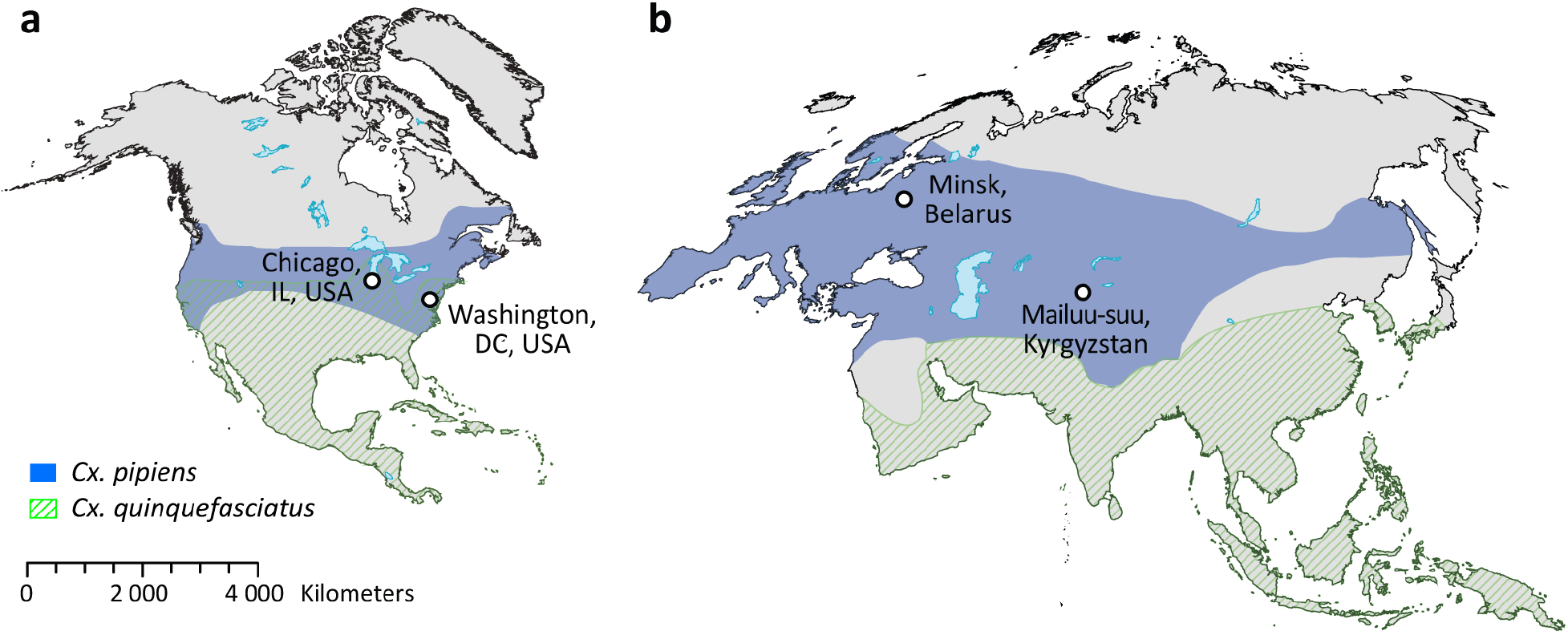
Mosquito collection sites in North America (a) and Eurasia (b). Worldwide distribution of *Cx. pipiens* and *Cx. quinquefasciatus* is indicated by blue color and green lines. The overlap between the sheds represents a species hybrid zone indicating that collections in the USA (Chicago, IL and Washington, D.C.) were made within the hybrid zone.

### Nuclear genome analysis

After the resequencing the 40 individual genomes, we performed neighbor joining analysis. The *Cx. quinquefasciatus* genome assembly ^24,28^ was used as a reference genome and an outgroup. The analysis identified two major monophyletic clusters that segregated in a subspecies-specific manner with high bootstrap support (Fig. 2). *Cx. p. molestus* and *Cx. p. pipiens* clusters included field collected mosquitoes from the Republic of Belarus and the Kyrgyz Republic as well as colonies from Chicago, IL, USA. In addition, we determined the presence of a third cluster that had a polyphyletic nature and included mosquitoes from *Cx. pipiens* autogenic and *Cx. pipiens* anautogenic colonies from Washington, D.C., USA and *Cx. quinquefasciatus*, based on the genome assembly of this species ^24,28^. In fact, the Washington, DC area is located within the *Cx. pipiens* - *Cx. quinqufasciatus* hybrid zone (Farajiollahi, et al. 2011) and, thus, these mosquitoes may represent hybrids between the latter species not included in the analysis.

**Fig. 2.**
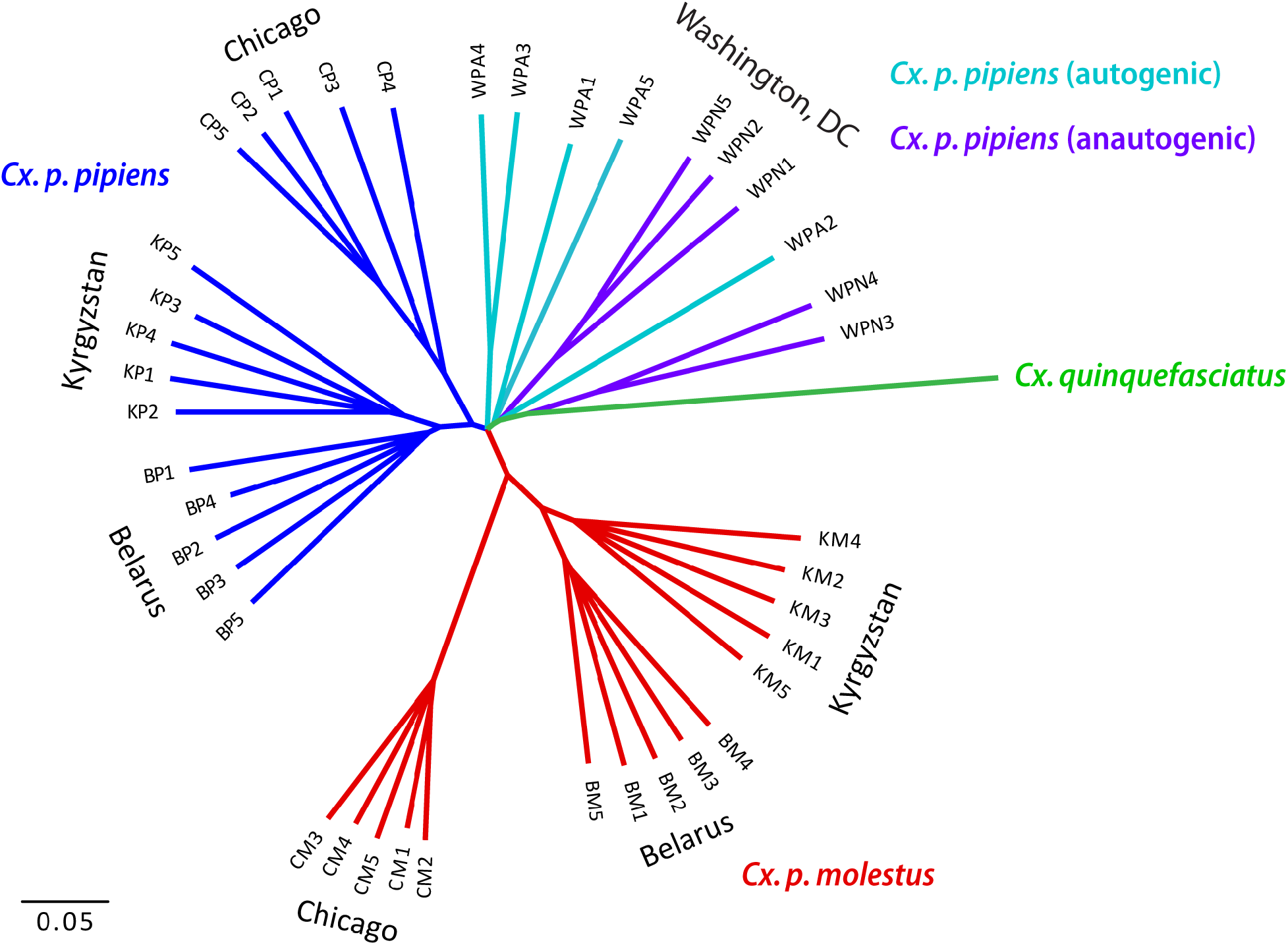
Neighbor joining tree based on K2P distances and autosomal genome-wide SNVs. Samples of *Cx. p. pipiens* and *Cx. p. molestus* from the Republic of Belarus (Belarus), the Kyrgyz Republic (Kyrgyzstan), and the USA (Chicago, IL) demonstrate their monophyletic origin except for the autogenic (WPA) and anautogenic (WPN) samples from Washington, D.C., USA. *Cx. quinquefasciatus* is used for the outgroup species. All the nodes with bootstrap support less than 90% were collapsed.

PCA analyses demonstrated groupings along the first principal component (13.57% of variance explained) with a closer relationship among *Cx. p. molestus* and *Cx. p. pipiens* subspecies rather than between geographical groups (Fig. 3). The second and third principal components (11.65% and 7.53%, respectively) grouped mainly geographical populations. In this analysis, both autogenic and anautogenic mosquitoes from Washington, D.C. grouped together with *Cx. p. pipiens* from other locations. The ADMIXTURE analysis demonstrated that *Cx. p. molestus* and *Cx. p. pipiens* from the Republic of Belarus, the Kyrgyz Republic, and Chicago, IL, USA formed subspecies-specific clusters at K=2, demonstrating the major pattern of population differentiation (Fig. 4). The specimens from Washington, DC clustered together with *Cx. p. pipiens* at K=2-3. We observed some putative signatures of admixture at K=2-3 in all *Cx. p. pipens* samples from the Republic of Belarus, the Kyrgyz Republic, in 2 out of 5 individuals in the *Cx. p. pipens* Chicago colony, and in all autogenic and 4 out of 5 anautogenic mosquitoes from Washington, D.C. Overall, the introgression signature between the subspecies was lower in northern populations of *Cx. p. pipiens* in the Republic of Belarus and Chicago, IL than in southern locations in the Kyrgyz Republic and Washington, D.C. In the Washington, D.C. samples, the admixture levels were higher in autogenic strains overall than in anautogenic strains. Interestingly, the signature of admixture in all *Cx. p. molestus* samples was very low or even absent, suggesting restricted gene flow from *Cx. p. pipiens* to *Cx. p. molestus*. At a higher level of clustering (K=6-9), the local populations of the subspecies formed distant well-defined clusters without high overlap, especially for *Cx. p. molestus*. Thus, we concluded that both autogenic and anautogenic samples from Washington, D.C. represent *Cx. p. pipiens* or *Cx. p. pipiens* - *Cx. quinqufasciatus* hybrids with some admixture of *Cx. p. molestus*, which is more pronounced in autogenic samples.

**Fig. 3.**
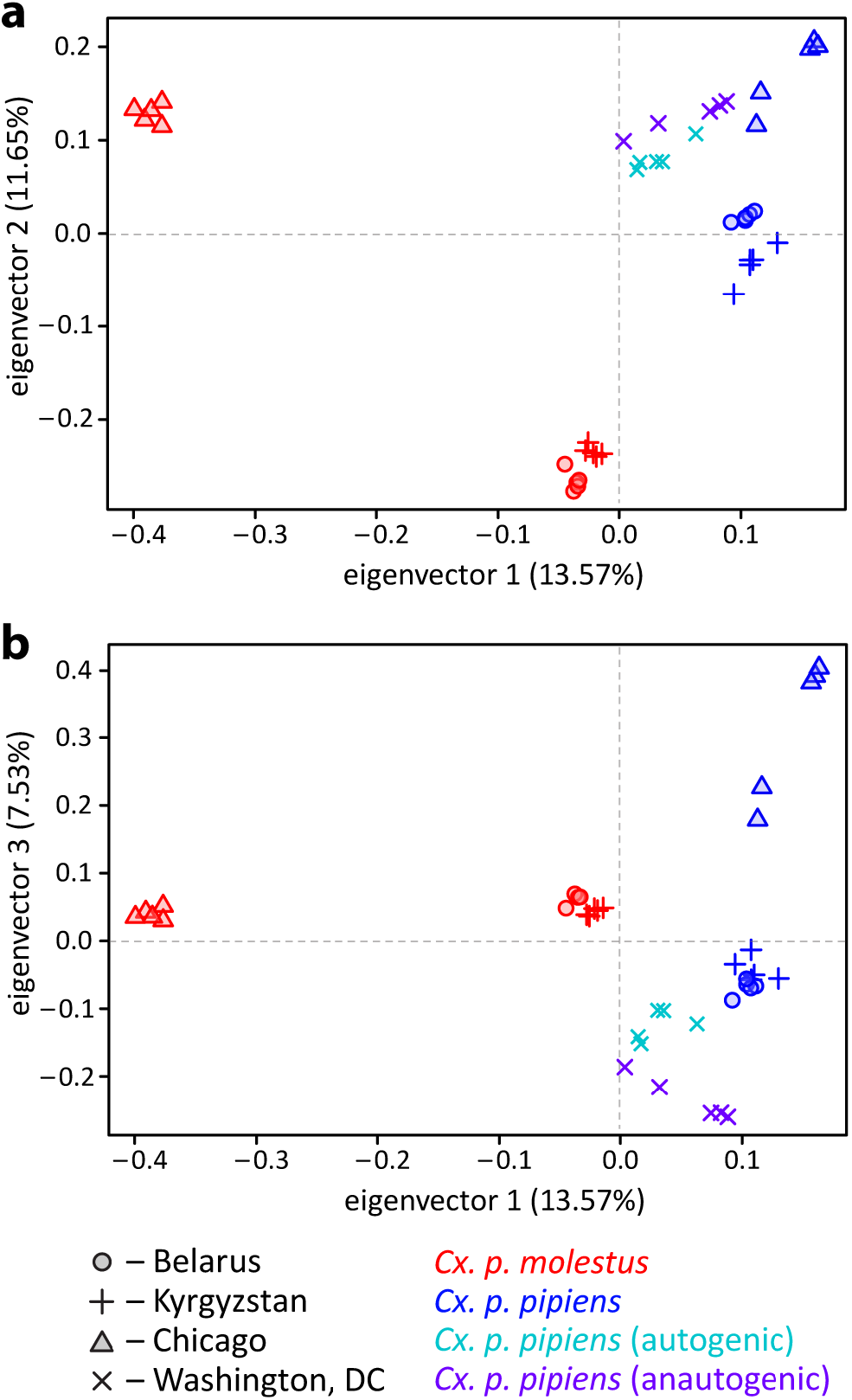
Principal component analysis of the individual samples for PC1-2 (a) and PC1-3 (b). PC1 separates the subspecies *Cx. p. pipiens* and *Cx. p. molestus* and PC2 with PC3 into separate geographic regions. Samples are from the Washington, D.C., USA group with other samples of *Cx. p. pipiens*.

**Fig. 4.**
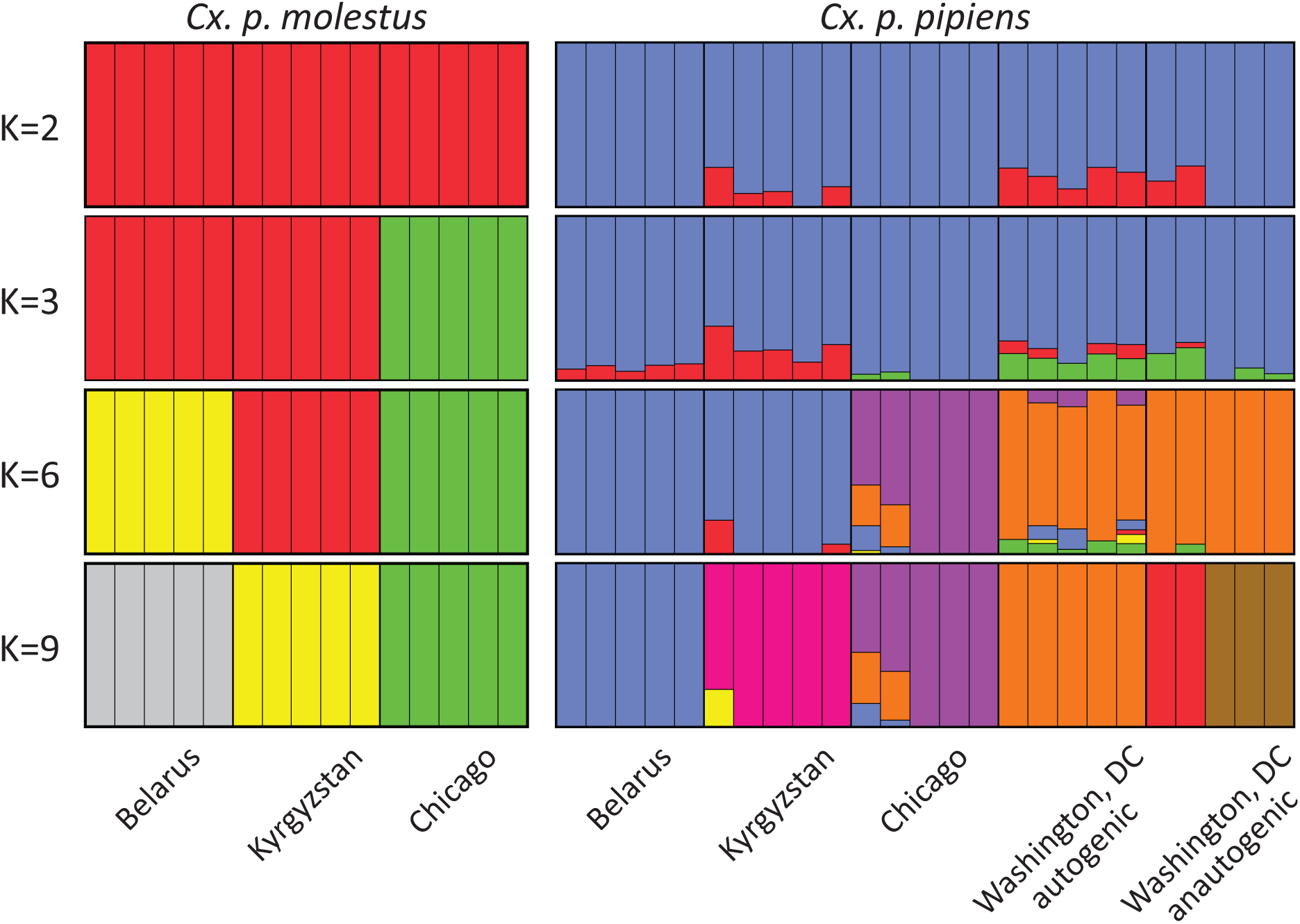
ADMIXTURE plot of the samples grouped by subspecies and regions. The major pattern of clustering is formed by subspecies level at K=2-3 and local populations at K=6-9. Samples are from the Washington, D.C., USA group with other samples of *Cx. p. pipiens*.

To get more insight into the nature of genomic differentiation between subspecies/populations, we calculated genome-wide pairwise *Fst* values and found that they varied between 0.08 and 0.25 and were highly significant for all comparisons (Fig. 5). The most diverged group was *Cx. p. molestus* from the Chicago strain, which can be explained by the very low genetic diversity of this strain caused by its longtime cultivation and possible bottleneck, which accelerated the genetic drift (Fig. 7). *Fst* analysis along the chromosomes demonstrated relatively uniform patterns of genetic differentiation across the genome without clear signs of “islands of divergence” (Fig. 6), which can indicate infrequent hybridization in nature and/or prezygotic instead of postzygotic reproductive isolation as major mechanisms of the subspecies differentiation. The *Fst* levels dropped around centromeres probably because a small number of markers are present in these regions due to a highly repetitive sequence composition. Overall, *Fst* values were higher between the subspecies than between autogenic and anautogenic *Cx. p. pipiens* from Washington, D.C. The level of genomic diversity did not differ significantly between the subspecies demonstrating an overall trend of slightly lower genetic diversity in *Cx. p. molestus;* however, this was not significant in pairwise comparisons except for the cultivated colonies from Chicago, IL (Fig. 7), which demonstrated a strong depletion of genetic diversity in *Cx. p. molestus*.

**Fig. 5.**
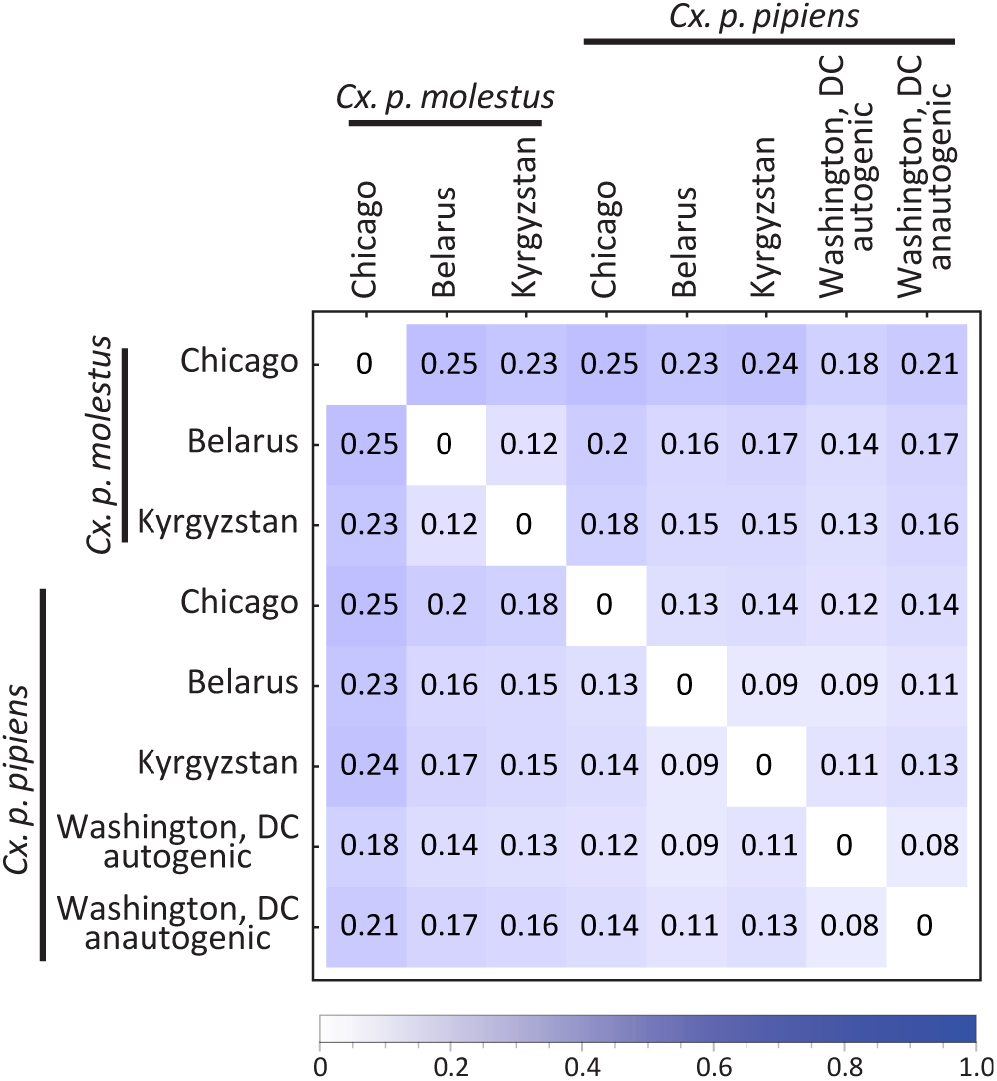
*Fst* matrix between the studied species/locations. *Cx. p. molestus* is the most divergent group due to its high level of genetic drift and low diversity. All values are highly significant.

**Fig. 6.**
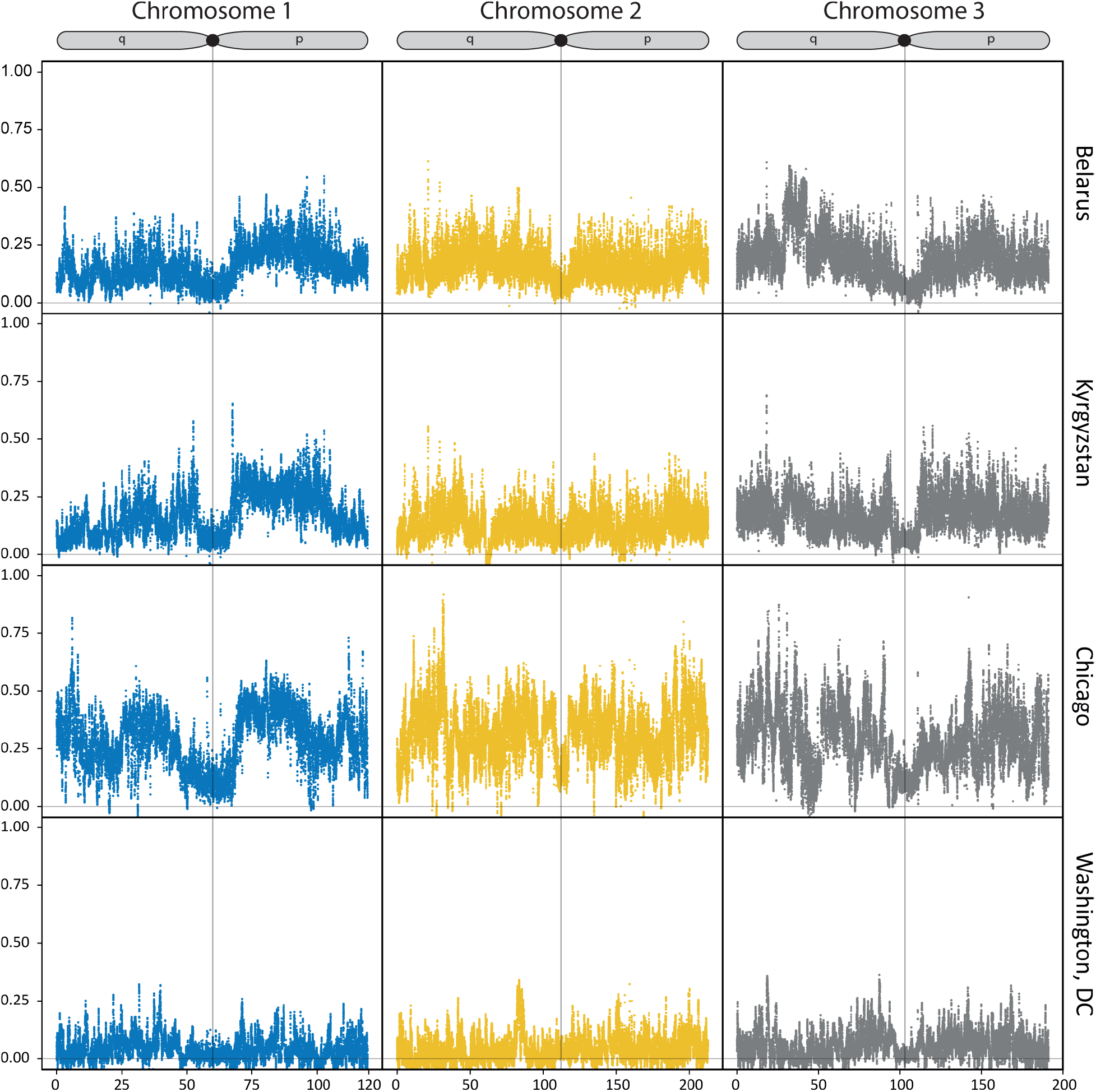
*Fst* values between the subspecies in different locations plotted along chromosomes. The patterns of genetic differentiation across the genome are more or less uniform. Overall, *Fst* values are higher between the subspecies than between autogenic and anautogenic *Cx. p. pipiens* from Washington, D.C., USA. P and q stand for short and long chromosome arms, respectively.

**Fig. 7.**
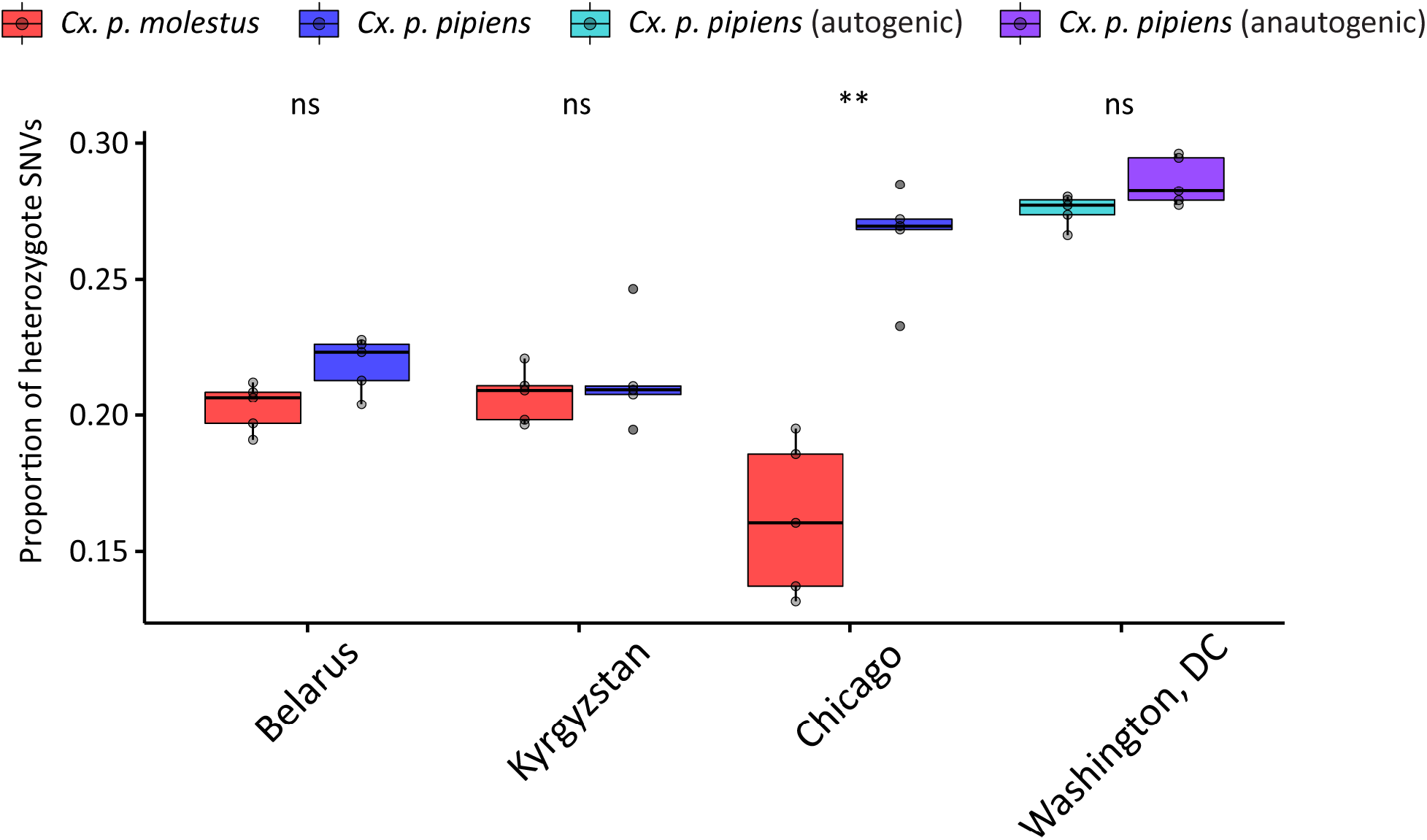
Level of individual genetic diversity estimated as a proportion of heterozygous SNVs per genome. Samples of *Cx. p. molestus* from Chicago, IL, USA demonstrate the lowest level of diversity and are significantly different from *Cx. p. pipiens* from the same location. ** indicate significant differences (P-value < 0.01).

### Mitochondrial genome analysis

In addition, to nuclear genome comparisons, we performed neighboring joining tree analyses using almost complete mitochondrial genomes for all 40 specimens recovered from the whole genome sequencing data. The patterns of phylogenetic structure were very different from the nuclear counterpart (Fig. 8) and showed the paraphyletic origin of the major haplogroups among the subspecies but the monophyletic structure between the continents. The mitochondrial genome of *Cx. quinquefasciatus* grouped inside the American haplogroup along with the samples from Washington, D.C. and did not demonstrate any significant divergence. Samples of *Cx. p. molestus* from the Republic of Belarus and the Kyrgyz Republic formed a diverged monophyletic haplogroup. Interestingly, two samples of *Cx. p. pipiens* from the Republic of Belarus represented a very diverged haplogroup, which may be reminiscent of the past species diversity or has been introgressed from other diverged populations/species. We specifically checked the quality of alignment and calling for this haplogroup but did not identify any abnormalities. Overall, we concluded that mitochondrial genome comparison does not reflect the true evolutionary history of the subspecies/species, probably due to multiple introgression events.

**Fig. 8.**
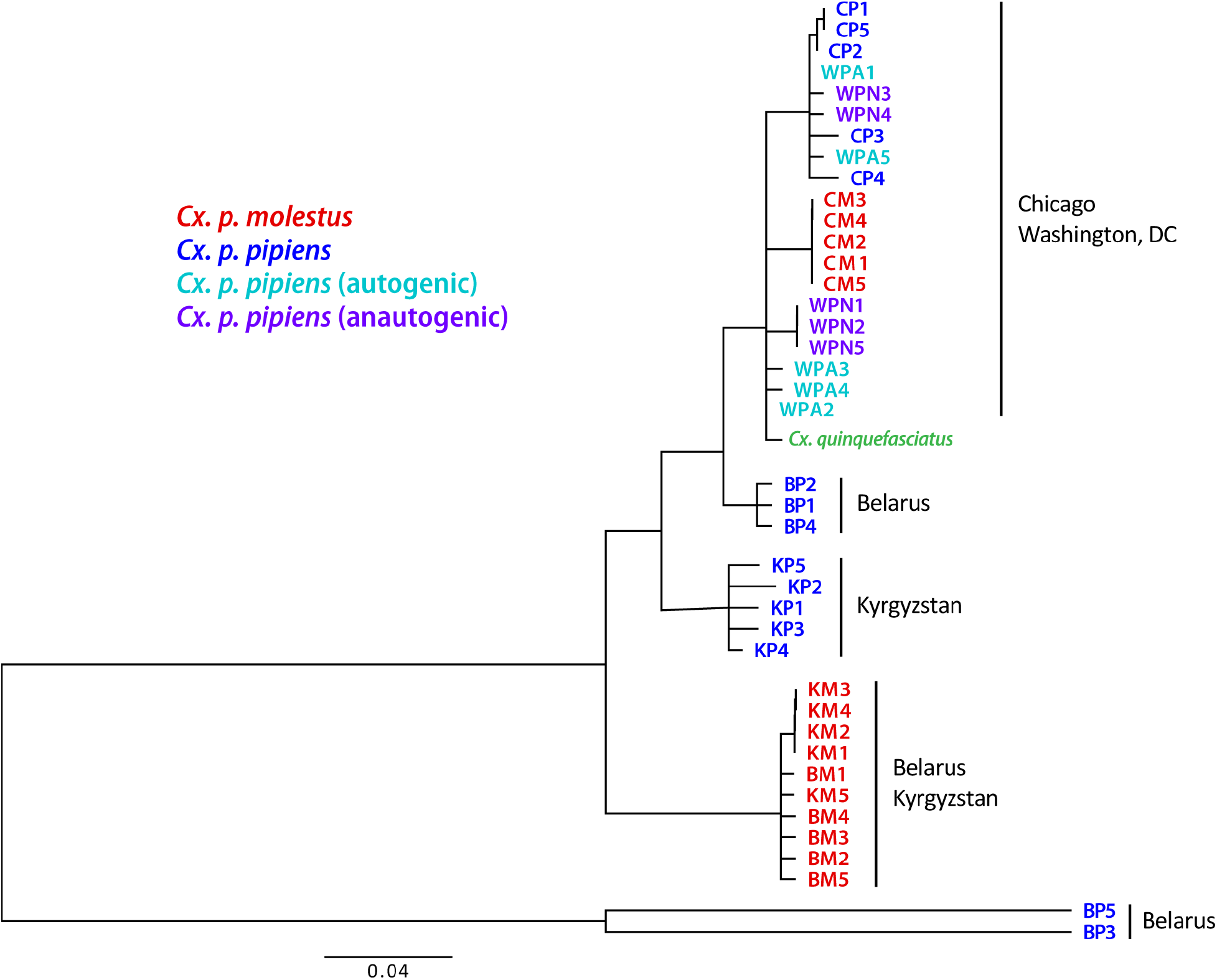
Neighbor joining tree based on the almost complete mtDNA genomes derived from WGS data (K2P distance). Patterns showed paraphyletic origins of the major haplogroups among the subspecies but monophyletic structures between the continents. All the nodes with bootstraps lower than 70% were collapsed.

## Discussion

### Independent monophyletic origin of *Culex pipiens pipiens* and *Culex pipiens molestus*

Two alternative hypotheses have been proposed to explain the differences in ecological and physiological strategies of *Cx. p. pipiens* and *Cx. p. molestus* ^18^. One hypothesis explains such differentiation as a rapid shift in physiological and behavior traits as an adaptation to an underground environment that is associated with human activities ^21^. In this scenario, the local populations of *Cx. p. molestus* must be closely related to the local populations of *Cx. p. pipiens* ^29^. This scenario considers *Cx. p. molestus* as an eco-physiological variant of *Cx. p. pipiens*. An alternative hypothesis suggests that the difference between *Cx. p. pipiens* and *Cx. p. molestus* is a result of the distinction in their evolutionary history. Ecological and physiological strategies of *Cx. p. molestus* could have first arisen under the warm climate ^18^. Later, such a strategy appeared to be relevant because of the human activities that created an underground environment and, thus, spread *Cx. p. molestus* all over the world. This scenario considers *Cx. p. molestus* and *Cx. p. pipiens* as separate evolutionary entities.

In our study, we tested these hypotheses using whole genome analysis of 40 *Cx. p. molestus* and *Cx. p. pipiens* samples collected from four locations in Eurasia and North America. Our findings rejected the hypothesis of reemergence of *Cx. p. molestus* from the local populations of *Cx. p. pipiens* and strongly supported the idea of an independent monophyletic origin of both *Cx. p. pipiens* and *Cx. p. molestus* from different continents. Neighbor joining analyses separately cluster *Cx. p. pipiens* and *Cx. p. molestus* from the Republic of Belarus, the Kyrgyz Republic, and Chicago, IL, USA (Fig. 2). The presence of an additional hybrid cluster formed by autogenic and anautogenic field collected specimens from Washington, D.C., USA more likely reflects the existence of a *Cx. pipiens* – *Cx. quinquefasciatus* hybrid zone in this area ^15^. However, based on genetic distances, this cluster is much closer related to the Eurasian *Cx. p. pipiens* rather than to the *Cx. p. molestus* from Eurasia and the USA (Chicago, IL). Additionally, PCA (Fig. 3) and ADMIXTURE (Fig. 4) analyses identified two distinct genetic clusters that correspond to *Cx. p. pipiens* and *Cx. p. molestus*. In both cases, all the specimens from Washington, D.C. clustered together with *Cx. p. pipiens* from other locations. Genome-wide pairwise *Fst* values were highly significant for all the comparisons (Fig. 5). We did not identify any specific region of high divergence in the genome (Fig. 6). Overall, *Fst* values were much higher between *Cx. p. molestus* and *Cx. p. pipiens* than between autogenic and anautogenic samples of *Cx. p. pipiens* from Washington, D.C. The most diverged group, based on all conducted analyses, was the strain of *Cx. p. molestus* from Chicago, IL, which can be explained by the very low genetic diversity of this strain caused by its longtime colonization and possible bottleneck, which accelerated genetic drift (Fig. 7).

Similar observations about the independent monophyletic origin of *Cx. p. molestus* and *Cx. p. pipiens* were obtained using microsatellite analysis in ~600 samples of *Cx. pipiens* mosquitoes derived from different world-wide locations ^18^. The unrooted distance tree procedure clustered aboveground populations of *Cx. p. pipiens* and underground populations of *Cx. p. molestus* from northern and southern Europe separately. Similar to our study, the aboveground populations of *Cx. p. pipiens* from the USA formed an additional cluster suggesting more intense hybridization between the members of the *Cx. pipiens* complex in North America. The admixture analysis indicated the presence of three major clusters that corresponded to *Cx. p. molestus, Cx. p. pipiens*, and *Cx. quinquefasciatus.* The latter cluster was found in this study only in the USA samples. Another work based on amplified fragment length polymorphism (AFLP) analyses compared samples from southern and northern Europe and strains from Chicago, IL, USA ^23^. The North American and European populations used in this study showed a similar ADMIXTURE pattern in the AFLP genome scan. The analysis of COI genes also indicated a monophyletic origin of *Cx. p. pipiens* and *Cx. p. molestus* in Europe, Asia, and Africa ^25^.

Thus, the independent monophyletic origin and high level of genetic divergence between *Cx. p. molestus* and *Cx. p. pipiens* suggest that these two members of the *Cx. pipiens* complex represent distinct phylogenetic entities with independent evolutionary histories prior to human-mediated translocation.

### Concepts of speciation and evolution of the *Culex pipiens* complex

Theoretical models of speciation in animals could be subdivided into two major groups: allopatric or geographical speciation and sympatric or ecological speciation. The first concept of speciation, which was intensively promoted by E. Mayr ^3^, suggests that incipient taxa first become isolated geographically. This situation reduces the gene flow between the populations and may lead to accumulation of mutations that cause genetic incompatibility among the hybrids. An alternative concept of speciation emphasizes ecological barriers between the emerging taxa as major drivers of evolution ^30^. This scenario considers the development of reproductive isolation between the populations as a result of adaptation to different environments without geographical isolation, which usually takes place in the face of gene flow. In this situation, the hybrids between incipient taxa become less fit to the environment that promotes natural selection of any traits that reduce mating between them. We think that overall diversification of the *Cx. p. pipiens* and *Cx. p. molestus* subspecies represents a striking example of speciation through isolation-by-ecology mechanisms. In our study, the *Fst* analysis determined significant levels of genomic divergence between *Cx. p. pipiens* and *Cx. p. molestus* (Fig. 5 and 6) across the entire genome without clear islands of speciation. Surprisingly, the levels of differentiation were lower around the centromeres, probably due to the low number of reliable SNVs in these highly repetitive regions. The differentiation was extremely high between the Chicago, IL, USA strains of *Cx. p. pipiens* and *Cx. p. molestus*, but was lower between the subspecies in the Eurasian mosquito collections. Overall, these observations suggest significant restriction of gene flow between the subspecies.

Several mechanisms of reproductive isolation between *Cx. p. pipiens* and *Cx. p. molestus* have been described ^16^. Two of them are prezygotic acting, before fertilization, and reduce the opportunities for mosquito mating. The first mechanism is related to habitat specialization of the larvae: *Cx. p. molestus* occupies basements or other underground environments, but *Cx. p. pipiens* prefers open aboveground water bodies for breeding sites. This reduces chances for the two subspecies to meet and mate at the adult stages. The second isolating mechanism relies on the differences in mating behavior between *Cx. p. molestus* and *Cx. p. pipiens*. *Cx. p. molestus* males usually form homogeneous swarms near the ground and require limited space for mating ^16^. In contrast, *Cx. p. pipiens* males swarm near the foliage about 2-3 m above the ground. Experimental studies of mating behavior in small cages indicate that in crosses of females and males of *Cx. p. molestus* copulation success was 90% but in *Cx. p. pipiens* it was only 3.3% ^31^. In inter-subspecies crosses between *Cx. p. molestus* and *Cx. p. pipiens*, the copulation success was also low and varied between 6.6% to 10% depending on the sexes of the subspecies. This study demonstrated that females of both subspecies actively avoid copulation with males from an alternative subspecies. Moreover, *Cx. p. pipiens* females were unsuccessful in receiving sperm from *Cx. p. molestus* and, as a result, produced no eggs.

Two other reproductive isolating mechanisms described for *Cx. p. pipiens* and *Cx. p. molestus* are postzygotic, they act after the mating and result in decreased fitness of the hybrids. One of the mechanisms is related to the inheritance of diapause in hybrids of *Cx. p. molestus* and *Cx. p. pipiens* as a recessive trait ^8^. F1 hybrids and a significant portion of F2 hybrids are unable to develop diapause and cannot survive under winter conditions. This mechanism, perhaps, could explain the higher introgression levels in *Cx. p. pipiens* in southern locations ^18,22,23^. Finally, the members of the *Cx. pipiens* complex are exposed to cytoplasmic incompatability of hybrids infected with different strains the *rickettsial* parasite *Wolbachia pipientis*. Despite cytoplasmic introgression of this parasite through hybridization between the members of the *Cx. pipiens* complex ^32^, cytoplasmic incompatibility could significantly limit survival rates of the hybrids. For example, *W. pipientis* infection significantly reduced hybridization between *Cx. pipiens* and *Cx. quinquefasciatus* in South Africa ^33^. Another study demonstrated that in Eurasian populations *Cx. p. molestus* was only infected by one strain of *W. pipientis* but *Cx. p. pipiens* by two different strains ^34^. Moreover, that the specimens of *Cx. p. molestus* and *Cx. p. pipiens*, which we used in our study, were infected with the same strains of *W. pipientis* in the south of the Kyrgyz Republic but by different strains in the north of the Republic of Belarus. Thus, it may explain the differences in introgression from *Cx. p. molestus* to *Cx. p. pipiens* that was more pronounced in the Republic of Belarus than in the Kyrgyz Republic. In fact, an interesting example of infectious speciation was described in the South American *Drosophila paulistorum* complex. In this complex, six semi-species with overlapping geographic distribution became reproductively isolated as a result of premating and postmating isolation triggered by the *Wolbachiae* infection ^35^.

Recent genomic studies conducted on different organisms including *Drosophila simulans* ^36^, *Rhagoletis* fruit flies ^37,38^, and *Heliconius* butterflies ^39,40^ provide additional evidence that ecological speciation occurs in nature. The genomic patterns of speciation can be very different ^2^ ranging from small genomic islands of speciation ^39^ to significant levels of divergence across the entire genome. Widespread genomic divergence was identified between the incipient species *Anopheles gambiae* and *An. coluzzii* in the *An. gambiae* complex ^41^. These species were originally identified by differences in the structure of their ribosomal DNA as S and M forms ^42^, but later their taxonomic status was elevated to the species level ^43^. *An. gambiae* and *An. coluzzii* are believed to develop reproductive barriers in sympatry as a result of the differences in their ecological preferences ^44^. The *An. gambiae* larval stage is associated with small rain pools. In contrast, *An. coluzzi* exploits persistent water reservoirs associated with rice cultivation. Although, premating barriers between the species are incomplete ^45^, they developed differences in their swarming behavior ^46,47^ and divergent song types ^48^.

Thus, a high level of genome-wide divergence, a striking difference in adaptation to distinct ecological environments and evidence of prezygotic and postzygotic barriers for mating suggest that *Cx. p. pipiens* and *Cx. p. molestus* represent distinct ecological units that undergo incipient ecological speciation.

### Hybridization in the *Culex pipiens* complex

The most intriguing observation of the members of the *Cx. pipiens* complex is that, despite differences in ecology, physiology, behavior, and geographic distribution, they still can hybridize and produce viable progeny in nature, indicating that reproductive isolation between them is not complete. Our study also demonstrated significant hybridization events between the subspecies *Cx. p. molestus* and *Cx. p. pipiens.* Whole genome comparison indicated that most of the *Cx. p. pipiens* samples represent individuals with some level of introgression from *Cx. p. molestus* (Fig. 4). We observed high discrepancy between the nuclear and mitochondrial phylogenies (Fig. 2, 8), which indicates that, historically, the transmission of mitochondrial genomes can happen between the subspecies. At the same time, we did not observe haplogroups shared between the local populations of the subspecies. In the continents, all the mitochondrial phylogenies were strongly monophyletic, which points to male-mediated dispersal and hybridization. The admixture with *Cx. p. molestus* was higher in southern populations of *Cx. p. pipiens* in the Kyrgyz Republic and in Washington, D.C., USA, but lower in northern populations in the Republic of Belarus and Chicago, IL, USA. This could be related to an incapability of the hybrids to develop diapause in cold climates. For comparison, microsatellite analysis revealed a modest level of hybridization between *Cx. p. pipiens* and *Cx. p. molestus* of ~8% in the northern cities of the USA (Chicago, IL and New York, NY) ^29,49^. In southern Europe, where *Cx. p. pipiens* and *Cx. p. molestus* could both be found aboveground, the hybridization levels between them were similar and was estimated at 8-10 % ^50^. Much higher levels of hybridization were found in southern populations in eastern USA, where ~40% of all samples were identified as hybrids between *Cx. p. molestus* and *Cx. p. pipiens* ^18^. Thus, overall hybridization rates between the members of *Cx. pipiens* were higher in North America than in the Old World. For comparison, hybridization between cryptic species in the *An. gambiae* complex (*An. gambiae* and *An. coluzzi*) significantly varied between 1% in Mali ^45^ to >20% in Guinea Bissau ^51^, which is comparable to overall hybridization levels between *Cx. p. molestus* and *Cx. p. pipiens.*

Intriguingly, in our study, we did not determine a *Cx. p. pipiens* admixture signature in any *Cx. p. molestus* samples from the Republic of Belarus, the Kyrgyz Republic, and Chicago, IL, USA (Fig. 4). These findings demonstrate very limited or no gene flow from *Cx. p. pipiens* to *Cx. p. molestus*. Similar findings of asymmetric introgression from *Cx. p. molestus* to *Cx. p. pipiens* was shown by previous studies ^19,50^. The mechanism of asymmetric introgression is currently unknown. One hypothesis suggests that males of *Cx. p. molestus*, which can mate in confined spaces, can hybridize with both *Cx. p. molestus* and *Cx. p. pipiens* females. In contrast, *Cx. p. pipiens* males, which require space for swarming, have a higher disposition to mate with *Cx. p. pipiens* females ^50^. However, more population genetic and experimental studies are needed to explain this phenomenon.

Finally, our study demonstrated that *Cx. p. pipiens* can develop autogeny as a result of adaptive introgression of the genetic material from *Cx. p. molestus*. In the field collected samples from aboveground and underground environments in Washington, D.C., USA, we selected mosquitoes for autogeny. The underground mosquitoes were autogenic but the aboveground mosquitoes were anautogenic. However, mosquitoes from both autogenic and anautogenic colonies formed a single cluster when neighbor joining analysis was applied (Fig. 2). Moreover, PCA (Fig. 3) and ADMIXTURE (Fig. 4) approaches cluster these samples together with *Cx. p. pipiens* from other locations. Populations with mixed characteristics were found in Europe ^50^ and in the USA ^52^. In Portugal, an unusual pattern of blood feeding behavior on birds was also found in *Cx. p. molestus* ^53^.

Thus, the presence of ongoing hybridization between members of the *Cx. pipiens* complex suggests that the speciation process between them is not complete and postzygotic barriers of reproductive isolation are not fully formed. Overall, we believe that members of the *Cx. pipiens* complex represent a remarkable model for studying different aspects of geographical and ecological speciation in the face of ongoing gene flow between them and local adaptations to diverse environments.

## Materials and Methods

### Mosquito collections

In this study, we used field collected mosquitoes from Minsk, Republic of Belarus, Mailuu-suu, Kyrgyz Republic, and Washington, D.C., USA (Table 1, Fig. 1). Mosquitoes from the Republic of Belarus were collected from an urban environment in Minsk, the capital city of the Republic. *Cx. p. pipiens* mosquitoes were collected at the adult stage in the basement of a multi-floor building. *Cx. p. molestus* were collected at the larval stage by dipping on the ground floor of a clinical hospital. In the Kyrgyz Republic, all samples were collected in an urban environment at the larval stage by dipping. *Cx. p. pipiens* were found in an open water pool, while *Cx. p. molestus* were found in an underground water barrel. Mosquitoes were identified as *Cx. p. molestus* and *Cx. p. pipiens* by the COI assay ^25^. In Washington, D.C., USA mosquitoes were collected in an urban environment as egg rafts from an open water puddle and an underground parking lot. Mosquitoes from both locations were successfully colonized and fed on fish pellet food at the larval stage. Morphologically, mosquito larvae from both collection sites were identified as *Cx. pipiens*. Adult mosquitoes were kept at 26 °C and offered 10% sucrose solution. Mosquitoes from both sites were tested for autogeny. Individuals from the parking lot demonstrated the ability to lay eggs without blood feeding, while individuals from the open water reservoir did not. Anautogenic mosquitoes were fed on artificial membrane blood feeders 4–5 days after emerging. Five mosquitoes from the second generation of both colonies were selected for sequencing. Finally, we used laboratory colonies of *Cx. p. molestus* and *Cx. p. pipiens* from Chicago, IL, USA. These colonies were established from mosquitoes collected in the Chicago, IL area. *Cx. p. molestus* was collected by sampling a drainage sump using aspirators and larval dipping in January 2009 (Mutebi and Savage, 2009) and *Cx. p. pipiens* was collected as egg rafts in above ground breeding sites in Evanston and Northfield, IL in August of 2016.

### DNA extraction and sequencing

DNA was extracted from individual mosquitoes for all Eurasian samples using the DNA Invisorb Spin Tissue Mini Kit (Invitek, Berlin, Germany) and for all American samples using the Qiagen Blood and Tissue Kit (Qiagen, Germantown, MD, USA). DNA concentration was determined by nanodrop; ~50 ng of DNA from each individual mosquito was sent for sequencing to the Fasteris company (Fasteris Inc., Switzerland). Sequencing was done using Illumina HiSeq 2 x 150 single index configuration with a 12-17X coverage for individual mosquito samples.

### Genomic analysis: variant calling

Individual genomes for each population were aligned against the *Cx. quinquefasciatus* reference genome ^28^ using BWA-MEM software ^54^. The resulting BAM files were sorted and PCR duplicates were removed using Samtools ^55^. To call Single Nucleotide Variations (SNVs), we used the *bcftools mpileup* and *call* multisample (*-m*) functions of the bcftools package ^56^ and had alignments with the quality of mapping equal to at least 40 and a base quality not less than 20. All of the BAM files were called simultaneously. The raw VCF file was filtered with vcftools ^57^ to remove all variants with a quality less than 500 *(--minQ 500*), more than two alleles per position, and were monomorphic with missing genotypes in more than four individuals.

To call mitochondrial DNA (mtDNA) variants, the reads were aligned to the nearly complete mitochondrial genome of *Cx. quinquefasciatus* with BWA-MEM and only unique, properly paired alignments with the highest quality (60) were left for the sorting and deduplication steps. The mitochondrial variants were called using the bcftools consensus algorithm (*bcftools call -c)* ^56^.

### Population genetic analysis

To reduce the influence of the linkage between neighboring loci on population genetic analysis, we removed all loci located closer than 10 Kbp from each other using vcftools (*--thin 10000*). The resulting pruned dataset was used to compute the Principal Component Analysis (PCA) using SNPrelate software ^58^ and unsupervised clustering using ADMIXTURE ^59^ for K=2-9. Pairwise *Fst* values ^60^ between locations/subspecies were calculated with the Stacks package ^61^. To profile the *Fst* values between subspecies along the chromosomes, we used the sliding window *Fst* approach implemented in vcftools with a window size equal to 50 Kbp and steps equal to 5 Kbp.

To construct neighbor-joining trees for the autosomal and mtDNA datasets, we used RapidNJ software ^62^ with K2P distance ^63^ and 1000 bootstrap replications.

## Data Availability

Genomic sequencing data will be available via NCBI upon the publication of the manuscript.

## Acknowledgments

We thank Igor V. Sharakhov for the productive discussion of the manuscript, Dmitri A. Karagodin for the help with formatting figures and Janet Webster for proofreading the text.

## Author contribution

MVS and AAY designed the experiments. RAM, MLF, AKS, and NVK performed mosquito collections and species identification. AAY conducted statistical and bioinformatics analysis. MLF provided mosquito colonies. MVS, AAY, and RAM wrote the manuscript. All authors read and approved the manuscript.

